# Cultivation-based identification of microorganisms in metalworking fluids and their role in hydrocarbon degradation

**DOI:** 10.64898/2026.03.18.712622

**Authors:** Adrian Heckel, Berke Ovat, Jan Reichinger, Nico Hanenkamp, Andreas Burkovski

## Abstract

Water-miscible metalworking fluids are widely used in industrial processes. Despite the fact that they contain biocides, they are almost always colonized by microorganisms, which degrade different components of the liquid, may clog machines due to biofilm formation and might pose a health risk to workers. In this study, samples from four metalworking machines operated with the same metalworking concentrate from two different locations, were analyzed with respect to microbial growth. Twenty-seven bacterial species and one fungus were identified. From these, twenty species were not observed before as colonizers of metalworking fluids. Growth of microorganisms, resulting health risks, putative contamination pathways and metabolic pathways involved in biodegradation are analyzed and discussed in this study.

**Graphical Abstract:** 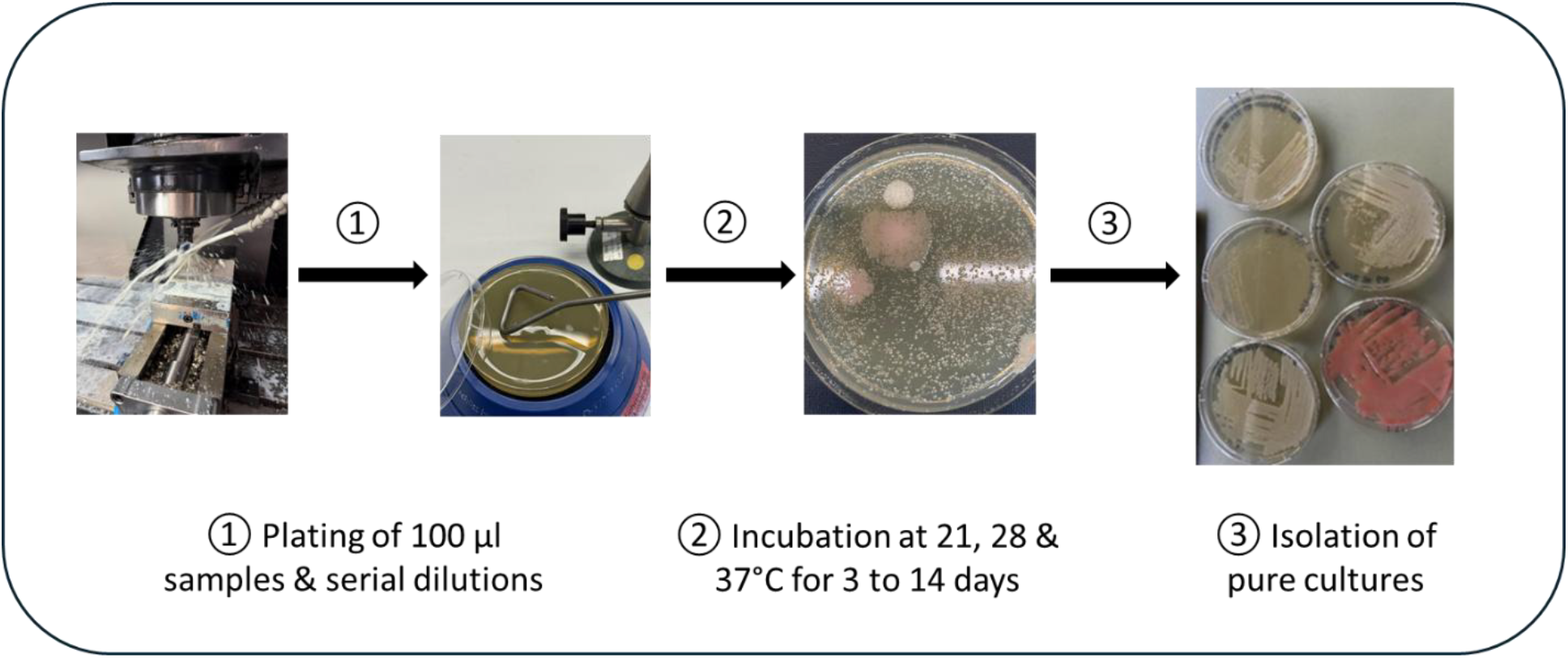

## 1. Introduction

Metalworking fluids (MWFs) are widely used in industry for cooling, lubrication, removal of metal chips and forming processes. In addition, MWFs support proper transformation of workpieces and protect tools from wearing down. Water-miscible MWFs, which account for approximately 90% of MWFs, typically consist of oil, water and application-specific additives, e.g. anti-corrosives, anti-foam agents and metal passivators (Brinksmeier et al., 2015; Passman & Küenzi, 2020). During use, MWFs are prone to microbial colonization (Mattsby-Baltzer et al., 1989; Sloyer et al., 2002), which is leading to severe detrimental effects. Biodeterioration of MWFs can change the stability of the water-oil emulsion, reduce viscosity and decrease pH, rendering the coolant and lubrication fluid dysfunctional, while fungal growth and highly resistant biofilms formed by microbes may cause clogging of machine parts and biofilm-based corrosion (Theaker & Thompson, 2010; Saha & Donofrio, 2012; Trafny, 2013; Passman & Küenzi, 2020). In addition to these technical problems, microbes may cause a considerable health risk to workers. Skin contact with contaminated MWFs may cause dermatitis (Grattan et al., 1989; Foulds & Koh, 1990; Alomar, 1994; Goh & Gan, 1994), while inhalation may lead to lung problems such as hypersensitivity pneumonitis or chronic obstructive pulmonary disease (Bernstein et al., 1995; Kreiss & Cox-Ganser, 1997; Shelton et al., 1999; Hodgson et al., 2001; Wilson et al., 2001; Fishwick et al., 2005; Rosenman, 2009; Tillie-Leblond et al., 2011; Burge, 2016).

Taken together, evaluation of bacterial contamination and control are crucial to optimize performance of MWFs and reduce health risks (Marchand et al., 2010). To monitor microbial growth, routinely dip slides (paddles) are used (Passman & Küenzi, 2020). While this method allows a semi-quantitative analysis of growth, it can only distinguish between bacteria and fungi. For deeper insight into the microbiology of MWFs a number of studies were carried out to identify microorganisms present in the MWF on species or genus level (for example, see van der Gast et al., 2003; Gilbert et al., 2009; for review, see Saha & Donofio, 2012). While often only a few species or genera are dominating MWF samples, the general microbiome composition seems to be highly variable (Lodders & Kämpfer, 2012). Main health risks observed in several studies seem to be members of the genus *Mycobacterium* (Shelton et al., 1999; Moore et al. 2000; Hodgson et al., 2001; Wilson et al., 2001; Wallace et al., 2002; Khan et al., 2005; Thorne et al., 2006).

Despite the industrial importance of MWFs and a number of studies carried out, our recent knowledge on the composition and function of microbes in MWFs is scarce. To add further detailed data on MWF-colonizing microorganisms and biofilm formation, we started a cultivation-based approach to identify members the microbiome of different machines operated with the same type of MWF and correlated the species level taxonomical information with available genome data.

## 2. Methods

### 2.1 Sample collection

Samples taken from four different machines at two locations, which were operated with the same MWF were analyzed in this study: a band saw, a lathe and two milling machines from the Chair of Resource and Energy-Efficient Production Machines (Fürth, Germany) and DMG Mori (Pfronten, Germany). The MWFs differed in terms of the degree of contamination and the area of application: Some machines were operated in three shifts five days a week and the concentration of the MWF was checked every two to three days and, if necessary, adjusted by adding water or concentrate to ensure stable conditions. Others were irregularly operated with extended downtimes for the MWF in the tank. The MWFs analyzed based on the mineral oil-based MWF concentrate ECO COOL GLOBAL 1000 (Fuchs GmbH, Mannheim, Germany).

### 2.2 Cultivation of samples and isolation of pure cultures

For the isolation of microbes 100 µl aliquots of MWF samples were spread in triplicate on following media: Columbia Blood Agar (Oxoid, Wesel, Germany), Luria Broth (LB) Agar (10 g tryptone, 5 g yeast extract, 10 g NaCl, 15 g agar per l), Meat Peptone Agar (3 g meat extract, 5 g peptone 15 g agar per l), Muller Hinton agar (Sigma, Steinheim, Germany), MWG agar (15 g agar, 10 ml Fuchs Ecocool Global 1000 per l), Sabouraud Agar (Merck, Darmstadt, Germany) and Tryptone Soy Agar (Roth, Karlsruhe, Germany). Subsequently, samples were incubated at 21, 28 and 37°C, respectively. Plates were incubated between three days and two weeks until considerable growth of colonies was detectable and pure cultures were isolated from these media.

To determine the growth properties of members of the core microbiome, corresponding type strains were purchased from the German Collection of Microorganisms and Cell Cultures (DSMZ, Braunschweig, Germany) and cultivated in LB medium (10 g tryptone, 5 g yeast extract, 10 g NaCl) under constant shaking at 28°C. All experiments were carried out in triplicate (independent biological replicates).

### 2.3 Identification of microorganisms

MALDI-ToF MS analysis was carried out by Labor Dr. Risch (Buchs, Switzerland) using fresh streak-outs of the isolates as this technique requires fresh biological material that contains proteins that have not yet undergone significant degradation (Reeve & Buddie, 2018; Topic Popovic et al., 2023). MALDI scores >2.0 are considered as accurate identification on species level, values between 1.7 and 2.0 represent moderate identification confidence and <1.7 no identification (Croxatto et al., 2012).

### 2.4 Analysis of metabolic pathways

To screen for metabolic pathways involved in hydrocarbon degradation, the Kyoto Encyclopedia of Genes and Genomes (KEGG, last assessed 16.03.2026) was screened for alkane degradation and fatty acid metabolism (pathway map “map00071”; Parekh et al., 1977).

## 3. Results

### 3.1 Growth characteristics of the MWF core microbiome

Based on the literature, members of the MWF core microbiome, the microorganisms most abundant in the different studies, are *Brevundimonas diminuta, Comamonas testosteroni, Morganella morganii, Mycobacterium immunogenum, Pseudomonas aeruginosa, Pseudomonas fluorescens, Pseudomonas putida, Pseudomonas oleovorans, Pseudomonas stutzeri, Shewanella putrefaciens* and *Stenotrophomonas maltophilia* (Cook & Gaylarde, 1988; Mattsby-Baltzer et al., 1989; Zacharisen et al., 1998; van der Gast et al., 2003; Veilette et al., 2004; Dilger et al., 2005; Khan et al., 2005; Cyprowski et al., 2007; Rabenstein et al., 2009; Gilbert et al., 2010; Marchand et al., 2010; Perkins & Angenent, 2010; Lodders & Kämpfer, 2012; Murat et al., 2012; Koch et al., 2015; Trafny et al., 2015; Kapoor et al., 2024). We suspected that one reason for their frequent abundance in MWFs may be a fast growth under standard laboratory conditions, making detection of the microorganisms easier. In fact, when cultivated in LB medium at 28°C under aeration these bacteria reached doubling times between 30 min and 230 min and an optical density at 600 nm (OD_600_) between 3.8 and 8.6 after 24 hours (Table 1).

**Tabelle 1.** Growth properties of the MWF core microbiome. Bacteria purchased from the German Collection of Microorganisms and Cell Cultures (DSMZ, Braunschweig, Germany) were grown in LB medium at 28°C under constant shaking. Experiments were carried out in triplicate (independent biological replicates and mean values ± standard deviations are shown.

| Species and strain | Doubling time [min] | OD<sub>600</sub> after 24 h |
| --- | --- | --- |
| *Brevundimonas diminuta* DSM 7234 | 162 ± 68 <sup>1)</sup> | 6.8 ± 0.4 |
| *Comamonas testosteroni* DSM 50244 | 30 ± 0 | 5.1 ± 0.2 |
| *Morganella morganii* DSM 14850 | 41 ± 2 | 4.9 ± 0.1 |
| *Mycobacterium immunogenum* DSM 45595 | 144 ± 1 | 4.0 ± 0.2 |
| *Pseudomonas aeruginosa* DSM 50071 | 58 ± 4 | 6.3 ± 0.2 |
| *Pseudomonas fluorescens* DSM 50090 | 34 ± 1 | 6.9 ± 0.1 |
| *Pseudomonas putida* DSM 291 | 43 ± 1 | 7.3 ± 0.4 |
| *Pseudomonas oleovorans* DSM 1045 | 101 ± 2 | 3.9 ± 0.1 |
| *Pseudomonas stutzeri* DSM 5190 | 95 ± 2 | 4.6 ± 0.6 |
| *Shewanella putrefaciens* DSM 6067 | 44 ± 2 | 5.4 ± 0.1 |
| *Stenotrophomonas maltophilia* DSM 50170 | 32 ± 1 | 8.2 ± 0.4 |

<sup>1)</sup> high standard deviation due to flocculating growth

### 3.2 Variability of microbial colonization of metalworking fluids

Samples were taken from four different MWFs and plated on the media described in chapter 2.2. To avoid a bias towards fast growing microorganisms and an overgrowth of slow growing microbes, serial dilutions were carried out and plates were incubated between three days and two weeks. Depending on the MWF and incubation conditions different microbial populations were observed (Figure 1).

**Figure 1.**
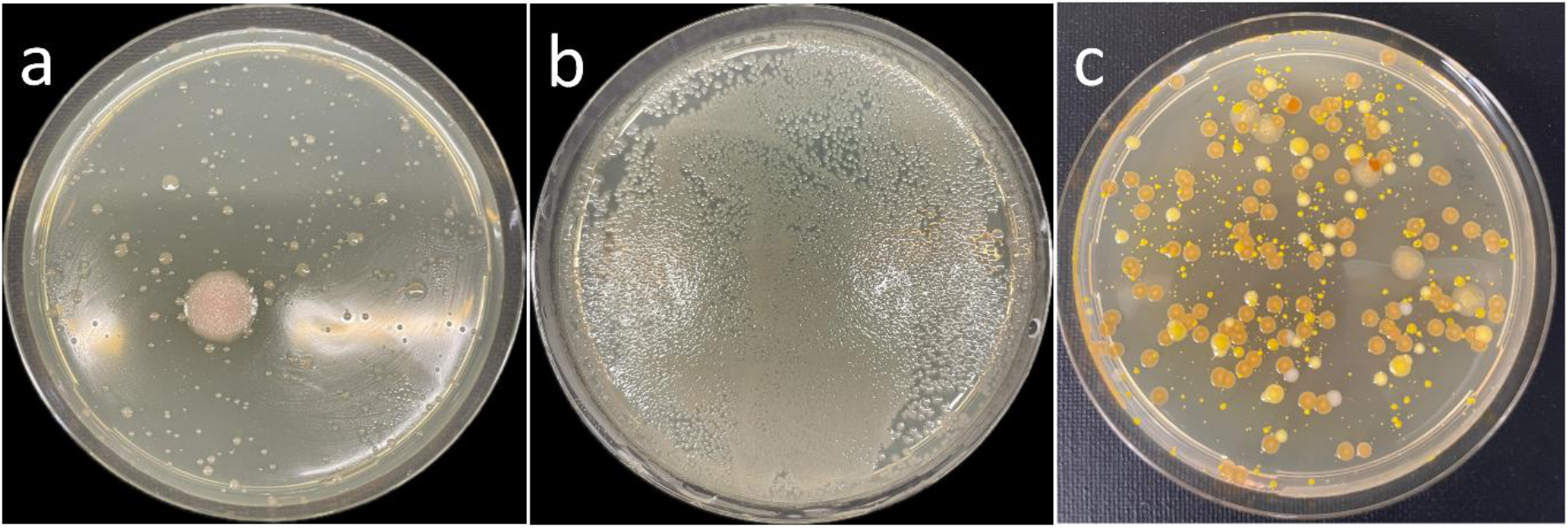
Examples of microorganisms isolated in this study. Streak-outs of (a), (b) MWFs, (c) water supply system.

In case of one of the machines, a band saw, massive biofilm formation was observed, which was analyzed further (Figure 2). Cultivation of the biofilm-forming microorganism and MALDI-ToF MS-assisted identification revealed that the leather-like structured biofilm was formed by *Scopulariopsis brevicaulis*, the arsenic fungus, a difficult-to-treat human pathogen causing onychomycosis (Mayser, 2025).

**Figure 2.**
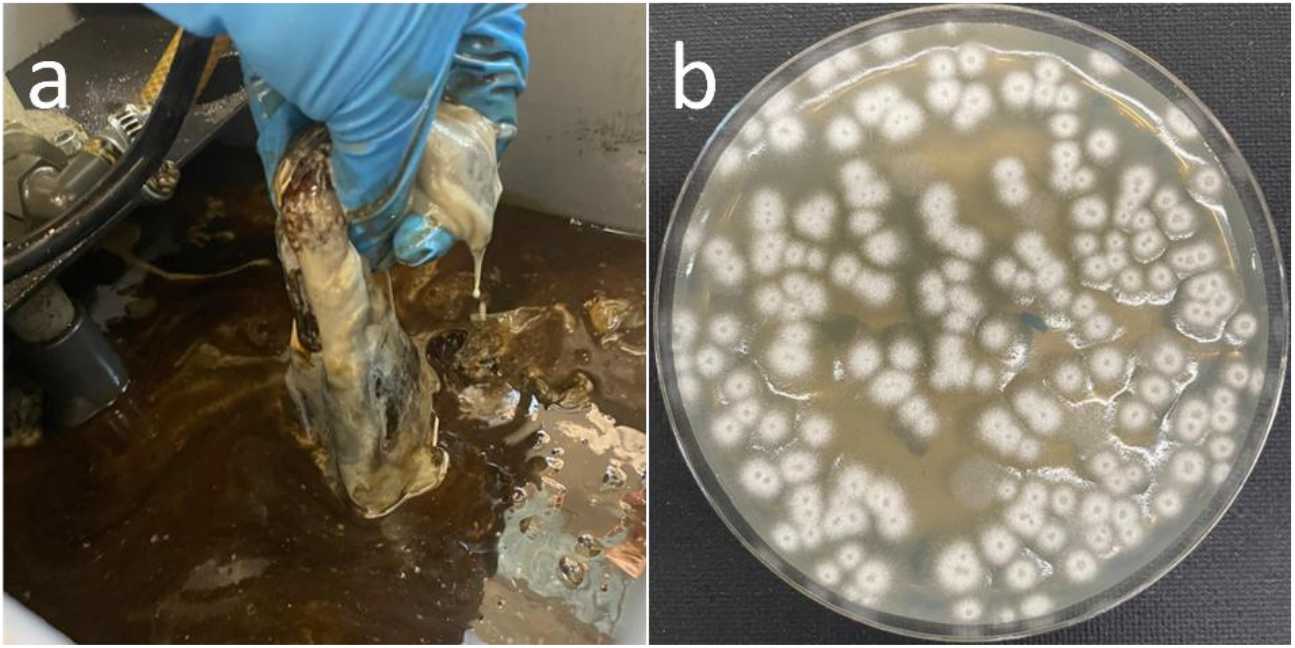
Biofilm formation in MWF. (a) Biofilm sample from band saw. (b) *Scopulariopsis brevicaulis* isolated from biofilm sample.

### 3.3 Identification of culturable microorganisms by MALDI-ToF MS

Microbial populations can be characterized by DNA extraction, 16S rRNA gene sequencing and subsequent taxonomical analysis. However, in case of MWFs, DNA extraction may be disturbed by components of the coolant (e.g. oil) and by contaminations from the workpieces (e.g. metal ions). Furthermore, identification by next generation sequencing often ends at family or genus level and the approach cannot distinguish between living and dead microbes. Therefore, we decided to concentrate on a culture-based approach based on the idea that microbes living in MWF should be rather robust and easy to handle. In fact, we were able to isolate twenty-seven different bacteria and one fungus on species level in this study (Table 2). To our best knowledge twenty of these were not found before in MWF. Twenty-two isolates could not be identified by MALDI-ToF-MS based on lacking databank entrances.

**Table 2.** Microorganisms identified in MWFs. Samples were cultivated under various conditions. Pure cultures were isolated and microorganisms subjected to identification via MALDI-ToF-MS. Species names, MALDI score and number of MWF samples with corresponding microorganism are given.

| Species | MALDI Score | Present in Samples |
| --- | --- | --- |
| *Arthrobacter globiformis* | 2.13 | 1 |
| *Bacillus pumilus* | 1.8-2.47 | 3 |
| *Comamonas sediminis* | 1.93 | 1 |

Table 2 cont.
|  |  |  |
| --- | --- | --- |
| *Corynebacterium accolens* | 2.01 | 1 |
| *Corynebacterium aurimucosum* | 2.02 | 1 |
| *Corynebacterium singulare* | 1.78 | 1 |
| *Corynebacterium tuberculostearicum* | 2.39 | 1 |
| *Cytobacillus horneckiae* | 2.39 | 1 |
| *Kocuria indica* | 1.91 | 1 |
| *Kocuria palustris* | 1.85 | 1 |
| *Kocuria rhizophila* | 1.93-2.08 | 2 |
| *Micrococcus luteus* | 1.83 | 1 |
| *Nocardiopsis dassonvillei* | 1.95 | 1 |
| *Paracoccus yeei* | 2.2 | 1 |
| *Priestia megaterium* (formerly *Bacillus megaterium*) | 2.14 | 1 |
| *Pseudomonas aeruginosa* | 2.38 | 1 |
| *Pseudomonas oleovorans* | 2.16-2.27 | 1 |
| *Scopulariopsis brevicaulis* | 2.45 | 1 |
| *Staphylococcus auricularis* | 1.91-1.92 | 2 |
| *Staphylococcus capitis* | 2.25-2.29 | 2 |
| *Staphylococcus epidermidis* | 1.98-2.17 | 3 |
| *Staphylococcus haemolyticus* | 2.29 | 1 |
| *Staphylococcus hominis* | 2.24-2.25 | 2 |
| *Staphylococcus nepalensis* | 1.77 | 1 |
| *Staphylococcus pragensis* | 2.45 | 1 |
| *Staphylococcus warneri* | 2.10 | 1 |
| *Streptomyces decoyicus* | 2.01 | 1 |
| *Streptomyces goshikiensis* | 2.34 | 1 |

When the microbiome of the MWFs from the different machines was analyzed with respect to the number of species identified, clear differences were observed. A maximum of eighteen species was found in one of the milling machines, while a mono-species biofilm was found in the MWF of the band saw analyzed in this study (Figure 3). As ECO COOL GLOBAL 1000 (Fuchs GmbH, Mannheim, Germany) was used as MWF concentrate in all machines, differences based on the MWF itself can be excluded and may be due to water used for MWF preparation, operation time and materials processed.

**Figure 3.**
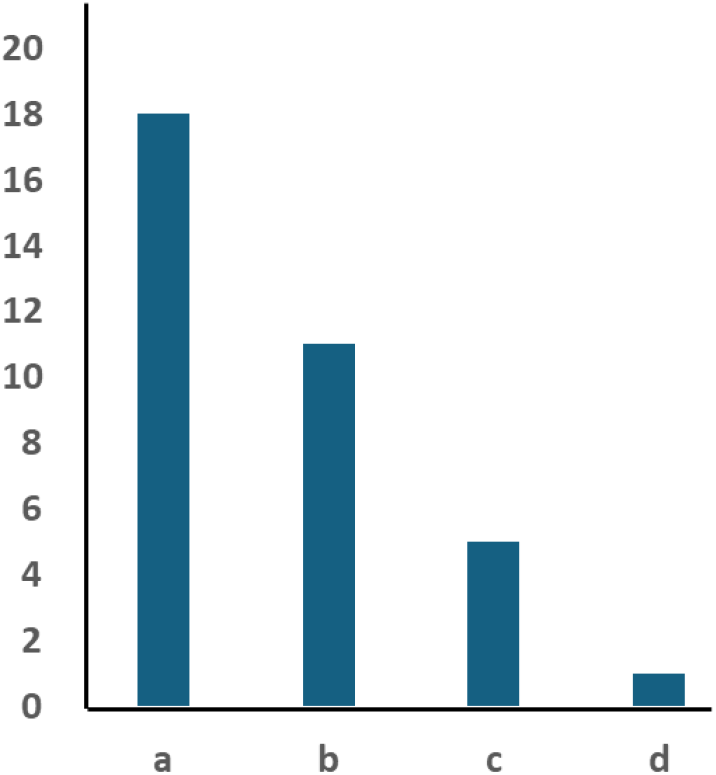
Species identified in MWFs of machines analyzed in this study. Columns indicate the number of identified species in MWF of the different machines. (a) milling machine and (b) lathe at the Chair of Resource and Energy-Efficient Production Machines, (c) milling machine at DMG Mori, (d) band saw at the Chair of Resource and Energy-Efficient Production Machines.

### 3.4 Isolation of human pathogens

As mentioned above, members of the MWF microbiome and in particular the genus *Mycobacterium*, may pose a risk to human health. Interestingly, in this study, mycobacteria were not identified as MWF colonizers. However, three groups of potentially harmful bacteria were identified, corynebacteria, pseudomonads and staphylococci.

From the four species of the genus *Corynebacterium* identified, three are at least occasional pathogens. *C. accolens* is in rare cases associated with inflammatory diseases such endocarditis, osteomyelitis and pneumonia (Claeys et al., 1994; Wong et al., 2010; Liu et al., 2021a; Liu et al.,2021b; Poucke et al., 2025). Interestingly, this species seems to have a protective effect against *Staphylococcus*.*aureus* and *Streptococcus pneumoniae* infections (Bomar et al., 2016, Menberu et al., 2021; Huang et al., 2022). *C. aurimucosum* was isolated from bone and joint infections, urinary tract infections, cases of bacteremia and skin and soft-tissue infections and is considered as an actual opportunistic pathogen (Lefevre et al., 2021). while *C. tuberculostearicum* is a human skin colonizer, which induces inflammatory signaling in skin cells and its relevance to chronic inflammatory diseases of the skin and cutaneous oncology is discussed (Altonsy et al., 2020).

While *P. aeruginosa* is a notorious nosocomial pathogen infecting cystic fibrosis patients (Nunez-Garcia et al., 2026) and causing severe wound, ocular and blood stream infections (Alharbi et al., 2025; Ather et al., 2025; Jabar et al., 2026), *P. oleovorans* is less in focus as a health risk. However, at least disinfectant-resistant biofilms formed by *P. oleovorans* were detected in endoscopes (Leeb-Zatorska et al., 2024) and a fatal case of *P. oleovorans* sepsis was reported (Gautam et al., 2015).

Many isolates of the genus *Staphylococcus* found in MWF, namely *S. auricularis, S. capitis, S. epidermidis, S. haemolyticus, S. hominis* and *S. warneri* are coagulase-negative staphylococci, which can cause skin and soft tissue infections, particularly in older and/or immunocompromised individuals (Natsis & Cohen, 2018). Epidemiologic and clinical studies have shown that *S. capitis, S. epidermidis, S. haemolyticus, S. hominis* and *S. warneri* may also be associated with meningitis (Azimi et al., 2020). In addition, *S. warneri* is known for its ability to colonize artificial prosthetic materials and its tropism for musculoskeletal and cardiovascular tissues (Ravaiolo et al., 2024).

### 3.5 Putative contamination pathways

Microorganisms can contaminate MWFs via planktonic cells in the water to prepare them or as part of biofilm consortia in pipes and water hoses used to add water to the MWFs. In fact, when testing the water supply system, a number of bacteria and a yeast were found (Table 3) including putative primary colonizers: *Acidovorax facilis* and *P. aeruginosa* are species exhibiting alkane and fatty acid degradation pathways, while *Pseudomonas putida* is able to degrade fatty acids by β-oxidation. Other putative water-borne contaminations found in this study are *C. sediminis* and *P. oleovorans*.

**Table 3.**
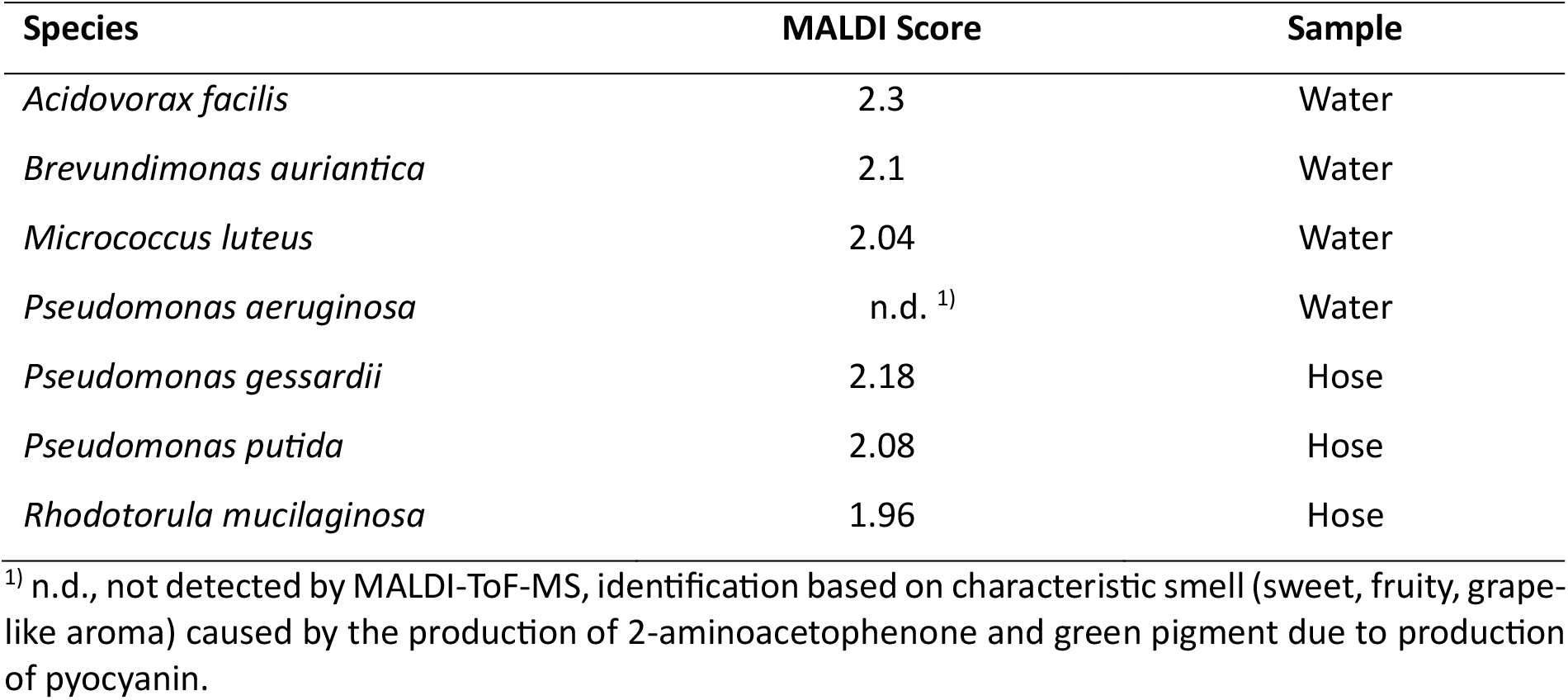
Microorganisms identified in water supply system. Samples were cultivated under various conditions. Pure cultures were isolated and microorganisms subjected to identification via MALDI-ToF-MS. Species names, MALDI score and sample sources are given.

| Species | MALDI Score | Sample |
| --- | --- | --- |
| *Acidovorax facilis* | 2.3 | Water |
| *Brevundimonas auriantica* | 2.1 | Water |
| *Micrococcus luteus* | 2.04 | Water |
| *Pseudomonas aeruginosa* | n.d. <sup>1)</sup> | Water |
| *Pseudomonas gessardii* | 2.18 | Hose |
| *Pseudomonas putida* | 2.08 | Hose |
| *Rhodotorula mucilaginosa* | 1.96 | Hose |

<sup>1)</sup> n.d., not detected by MALDI-ToF-MS, identification based on characteristic smell (sweet, fruity, grape-like aroma) caused by the production of 2-aminoacetophenone and green pigment due to production of pyocyanin.

Putative skin colonizers, especially corynebacteria and staphylococci, may be introduced in the MWF by the personnel handling machines, tools and metal pieces, while soil bacteria such as bacilli, streptomycetes, *Arthrobacter* and *Priestia* species may be introduced by dirt and dust and *M. luteus* is a typical air-borne contamination.

### 3.6 Influence of microbes on MWF quality

#### 3.6.1 Biosurfactant production

Microorganisms produce different classes of biosurfactants, surface-active agents that form micelles or vesicles around hydrocarbons to enhance uptake of hydrophobic substrates (Dini et al., 2024). In MWFs biosurfactants work as unwanted additional emulsifiers and support degradation of hydrocarbons. *P. aeruginosa* is well-known for its production of rhamnolipids (Das et al., 2025) and also *P. putida*, found in a biofilm sample of a hose in this study, naturally produce rhamnolipids although at a lesser extent (Toribio et al., 2010). In addition, glycolipid production was reported for *Arthobacter* strains (Campos et al., 2013; Santos et al., 2016).

#### 3.6.2 Hydrocarbon degradation pathways

In contrast to pure laboratory cultures, microbes form consortia in nature, which often work together to degrade complex nutrient sources. A well-investigated example in this regard is the microbial flora of the rumen (Weimer, 2022). In this part of the gastrointestinal tract of ruminants, cellulose and other polymers from plant material are metabolized and volatile fatty acids, carbon dioxide, hydrogen and methane are produced by bacteria and archaea.

As we suspected that similar interspecies cooperations may also play a role in MWF biodeterioration, we screened the KEGG data base for corresponding degradation pathways of the primary carbon sources in MWFs, which contain 1 to 5% petroleum oil and less than 0.1% fatty acids (Saha & Donofrio, 20212). Microorganisms convert alkanes via oxidation into fatty acids (van Beilen & Funhoff, 2007; Guo et al., 2026), which are then degraded by the β-oxidation pathway to acetyl-CoA and feed into the tricarboxylic acid cycle (TCA). Corresponding enzymes for alkane oxidation were found for example already in 1994 for *P. oleovorans* (van Beilen et al., 1994).

According to the KEGG database, only two species identified in this study as part of the MWF microbiome were able to degrade alkanes starting with an initial oxidation step of the carbon chain followed by β-oxidation. Members of this group of primary degraders were *P. yeei* and *P. aeruginosa*. For *Pseudomonas oleovorans*, an alkane oxidation pathway was not annotated in KEGG, but described before (van Beilen et al., 1994). Carbon utilization restricted to fatty acid consumption was found for *B. pumilus, C. sediminis, C. aurimucosum, K. palustris, P. megaterium, S. decoyicus* and *S. goshikiensis*. Growth of these microorganisms is only supported by alkanes, when these are oxidized and converted to fatty acids by the primary alkane degraders. Thirteen species identified here lacked any alkane and fatty acid degradation capacity, namely *C. accolens, C. singulare, K. indica, K. rhizophila, M. luteus*, N. *dassonvillei, S. auricularis, S. capitis, S. epidermidis, S. haemolyticus, S. hominis, S. nepalensis* and *S. warneri*. These bacteria will either degrade other components of the MWFs or metabolic products of alkane and fatty acid degraders. No data were provided by KEGG for *A. globiformis, C. tuberculostearicum, C. horneckiae, S. brevicaulis* and *S. pragensis* (Figure 4).

**Figure 4.**
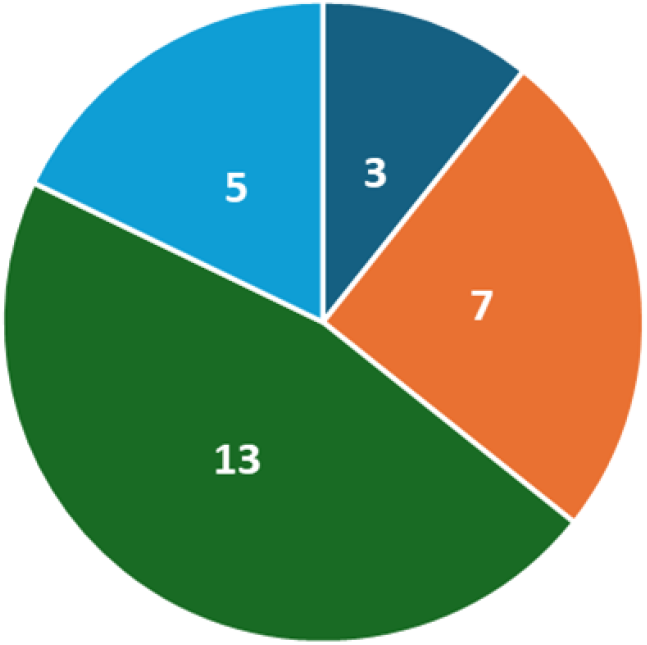
Pie chart of metabolic pathways involved in hydrocarbon degradation. Dark blue, number of species with alkane and β-oxidation pathway; orange, number of species with β-oxidation pathway; green, number of species without alkane and fatty acid degradation capacity; light blue, no genomic data available in KEGG.

## 4. Discussion

To our best knowledge, almost 100 microbial species were identified in MWFs up to now (Cook & Gaylarde, 1988; Mattsby-Baltzer et al., 1989; Zacharisen et al., 1998; van der Gast et al., 2003; Veilette et al., 2004; Dilger et al., 2005; Khan et al., 2005; Cyprowski et al., 2007; Rabenstein et al., 2009; Gilbert et al., 2010; Marchand et al., 2010; Perkins & Angenent, 2010; Lodders & Kämpfer, 2012; Murat et al., 2012; Koch et al., 2015; Trafny et al., 2015; Kapoor et al., 2024). In this study, 20 new species were identified by a cultivation-based approach adding further information on our knowledge on the MWF microbiome.

Based on the literature, *B. diminuta, C. testosteroni, M. morganii, M. immunogenum, P. aeruginosa, P. fluorescens, P. putida, P. oleovorans, P. stutzeri, S. putrefaciens* and *S. maltophilia* form the MWF core microbiome. While *Pseudomonas* species were frequently found in the different samples and also *Brevundimonas* and *Comamonas* species were identified, mycobacteria were not observed in this study. A putative reason for this result might be the fact that this study was based on the analysis of one distinct MWF, i.e. ECO COOL GLOBAL 1000 (Fuchs GmbH, Mannheim, Germany), instead of different ones as in other studies. Interestingly, even the tested samples of one distinct MWF showed a rather variable and individual colonization with a limited number of species as observed before for different MWFs (van der Gast et al., 2003; Gilbert et al., 2009; Lodder & Kämpfer, 2012).

The frequent colonization of MWFs with *Pseudomonas* species documented in literature and observed in this study, is not only detrimental for MWF quality and function, but may also pose a serious health risk to workers, which can be minimized with an optimized water supply. As described, tap water or water from wells and rivers may be used for MWF preparation (Lodder & Kämpfer, 2012). Reduction of water-borne microbial contaminations from these sources may be achieved by UV treatment (Barkhudarov et al., 2008; O’Flaherty et al., 2018), nano- and ultra-filtration processes (Schwermer et al., 2018) or ozone and hydrogen peroxide addition (Cho et al., 2010; Martirosyan et al., 2011). A further, very cost-effective approach may be the implementation of Boron-doped diamond electrodes for water sanitation (Schorr et al., 2019; Koch et al., 2022). These treatments may help to prolong MWF life cycle and reduce health risks for workers.

Future studies on MWFs may implement more quantitative analyses, e.g. real-time RT-PCR flanked by the determination of colony-forming units, and a time-resolved analysis of colonization, since our analyses of metabolic pathways hint a succession of microbes, starting with primary alkane degraders. In addition, resistance to high pH and microbial acid production might be important parameters influencing the composition of the MWF microbiome and MWF biodegradation.

## Abbreviations used in this study

KEGG: Kyoto Encyclopedia of Genes and Genomes
LB: Luria Broth
MALDI-ToF: MS Matrix-assisted laser desorption/ionization time of flight mass spectrometry
MWF: metalworking fluid
OD_600_: optical density at 600 nm
PBS: phosphate-buffered saline
RT PCR: reverse transcriptase polymerase chain reaction
TCA: tricarboxylic acid cycle
UV: ultraviolet

## Acknowledgements

The authors wish to thank Michael Kirbach (DMG Mori, Pfronten, Germany) for providing samples of used metalworking fluid.

## Author contributions

AH: investigation, methodology, formal analysis, visualization. BO: investigation, formal analysis, visualization. JR: project administration. NH: funding acquisition. AB: conceptualization, resources, writing—original draft preparation, writing—review and editing. All authors have read and agreed to the published version of the manuscript.

## Data availability

The raw data supporting the conclusions of this article will be made available by the authors on request.

## References

Alharbi, M.S., Moursi, S.A., Alshammari, A., Aboras, R., Rakha, E., Hossain, A., Alshubrumi, S., Alnazha, K., Khaja, A.S.S., Saleem, M. (2025) Multidrug-resistant Pseudomonas aeruginosa: Pathogenesis, resistance mechanisms, and novel therapeutic strategies. Virulence 16(1), 2580160. doi: 10.1080/21505594.2025.2580160.

Alomar, A. (1994) Occupational skin disease from cutting fluids. Dermatol. Clin. 12(3), 537–546.

Altonsy, M.O., Kurwa, H.A., Lauzon, G.J., Amrein, M., Gerber, A.N., Almishri, W., Mydlarski, P.R. (2020) Corynebacterium tuberculostearicum, a human skin colonizer, induces the canonical nuclear factor-κB inflammatory signaling pathway in human skin cells. Immun. Inflamm. Dis. 8(1), 62–79. doi: 10.1002/iid3.284.

Ather, M., Conrady, C.D. (2025) Emerging Ocular Pathogen Resistance and Clinically Used Solutions: A Problem That Is More than Meets the Eye. Pharmaceuticals (Basel) 19(1) 31. doi: 10.3390/ph19010031.

Azimi, T., Mirzadeh, M., Sabour, S., Nasser, A., Fallah, F., Pourmand, M.R. (2020) Coagulase-negative staphylococci (CoNS) meningitis: a narrative review of the literature from 2000 to 2020. New Microbes New Infect. 37, 100755. doi: 10.1016/j.nmni.2020.100755.

Barkhudarov, E.M., Christofi, N., Kossyi, I.A., Misakyan, M.A., Sharp, J., Taktakishvili, I.M. (2008) Killing bacteria present on surfaces in films or droplets using microwave UV lamps. World J. Microbiol. Biotechnol. 24, 761–769.

Bernstein, D.I., Lummus, Z.L., Santilli, G., Siskosky, J., Bernstein, I.L. (1995) Machine operator’s lung. A hypersensitivity pneumonitis disorder associated with exposure to metalworking fluid aerosols. Chest 108(3), 636–641. doi: 10.1378/chest.108.3.636.

Bomar, L., Brugger, S.D., Yost, B.H., Davies, S.S., Lemon, K.P. (2016) Corynebacterium accolens Releases Antipneumococcal Free Fatty Acids from Human Nostril and Skin Surface Triacylglycerols. mBio 7(1), e01725–15. doi: 10.1128/mBio.01725-15.

Brinksmeier, E., Meyer, D., Huesmann-Cordes, A.G., Herrmann, C. (2015) Metalworking fluids - Mechanisms and Performance. CIRP Annals – Manufact. Tech. 64, 605–628.

Burge, P.S. (2016) Hypersensitivity Pneumonitis Due to Metalworking Fluid Aerosols. Curr. Allergy Asthma Rep. 16(8), 59. doi: 10.1007/s11882-016-0639-0.

Campos, J.M., Stamford, T.L., Sarubbo, L.A., de Luna, J.M., Rufino, R.D., Banat, I.M. (2013) Microbial biosurfactants as additives for food industries. Biotechnol. Prog. 29(5), 1097–1108. doi: 10.1002/btpr.1796.

Cho, M., Kim, J., Kim, Y.J., Yoon, J., Kim, J.-H. (2010) Mechanisms of Escherichia coli inactivation by several disinfectants. Water Res. 44, 3410–3434.

Claeys, G., Vanhouteghem, H., Riegel, P., Wauters, G., Hamerlynck, R., Dierick, J., de Witte, J., Verschraegen, G., Vaneechoutte, M. (1996) Endocarditis of native aortic and mitral valves due to Corynebacterium accolens: report of a case and application of phenotypic and genotypic techniques for identification. J. Clin. Microbiol. 34(5), 1290–1292. doi: 10.1128/jcm.34.5.1290-1292.1996.

Cook, P.E., Gaylarde, C.C. (1988) Biofilm formation in aqueous metal working fluids. Int. Biodeterior. 24, 265–270. 10.1016/0265-3036(88)90010-3.

Croxatto, A., Prod’hom, G., Greub, G. (2012) Applications of MALDI-TOF mass spectrometry in clinical diagnostic microbiology. FEMS Microbiol. Rev. 36(2), 380–407. doi: 10.1111/j.1574-6976.2011.00298.x.

Cyprowski, M., Piotrowska, M., Zakowska, Z., Szadkowska-Stańczyk, I. (2007) Microbial and endotoxin contamination of water-soluble metalworking fluids. Int. J. Occup. Med. Environ. Health 20(4), 365–371. doi: 10.2478/v10001-007-0036-y.

Das, P.V., Valentine, M.E., Long, T.E., Yu, H.D. (2025) Attenuated Strains of Pseudomonas aeruginosa: A Promising Cell Factory for Rhamnolipid Production. Microb. Biotechnol. 18(11), e70239. doi: 10.1111/1751-7915.70239.

Dilger, S., Fluri, A., Sonntag, H.-G. (2005) Bacterial contamination of preserved and non-preserved metal working fluids. Int. J. Hyg. Environ. Health 208, 467.476. 10.1016/j.ijheh.2005.09.001.

Dini, S., Bekhit, A.E.A., Roohinejad, S., Vale, J.M., Agyei, D. (2024) The Physicochemical and Functional Properties of Biosurfactants: A Review. Molecules 29(11), 2544. doi: 10.3390/molecules29112544.

Fishwick, D., Tate, P., Elms, J., Robinson, E., Crook, B., Gallagher, F., Lennox, R., Curran, A. (2025) Respiratory symptoms, immunology and organism identification in contaminated metalworking fluid workers. What you see is not what you get. Occup. Med. 55(3), 238–241. doi: 10.1093/occmed/kqi049.

Foulds, I.S., Koh, D. (1990) Dermatitis from metalworking fluids. Clin. Exp. Dermatol. 15(3), 157–162. doi: 10.1111/j.1365-2230.1990.tb02062.x.

Gautam, L., Kaur, R., Kumar, S., Bansal, A., Gautam, V., Singh, M., Ray, P. 2015. Pseudomonas oleovorans Sepsis in a Child: The First Reported Case in India. Jpn. J. Infect. Dis. 68(3), 254–255. doi: 10.7883/yoken.JJID.2014.174.

Gilbert, Y., Veillette, M., Duchaine, C. (2010) Metalworking fluids biodiversity characterization. J. Appl. Microbiol. 108(2), 437–449. doi: 10.1111/j.1365-2672.2009.04433.x.

Goh, C.L., Gan, S.L. (1994) The incidence of cutting fluid dermatitis among metalworkers in a metal fabrication factory: a prospective study. Contact Dermatitis 31(2), 111–115. doi: 10.1111/j.1600-0536.1994.tb01929.x.

Grattan, C.E., English, J.S., Foulds, I.S., Rycroft, R.J. (1989) Cutting fluid dermatitis. Contact Dermatitis 20(5), 372–376. doi: 10.1111/j.1600-0536.1989.tb03175.x.

Guo, X., Zhu, S., Zhu, N., Zhang, S., Yang, S., Luo, G., Li, H., Wang, Y., Sun, J., Ma, B. (2026) Pan-Genomic and Phenotypic Characterisation of Petroleum Hydrocarbon Degradation by Pseudomonas Species. Environ. Microbiol. Rep. 18(1), e70300. doi: 10.1111/1758-2229.70300.

Hodgson, M.J., Bracker, A., Yang, C., Storey, E., Jarvis, B.J., Milton, D., Lummus, Z., Bernstein, D., Cole, S. (2001) Hypersensitivity pneumonitis in a metal-working environment. Am. J. Ind. Med. 39(6), 616–628. doi: 10.1002/ajim.1061.

Huang, S., Hon, K., Bennett, C., Hu, H., Menberu, M., Wormald, P.J., Zhao, Y., Vreugde, S., Liu, S. (2022) Corynebacterium accolens inhibits Staphylococcus aureus induced mucosal barrier disruption. Front. Microbiol. 13, 984741. doi: 10.3389/fmicb.2022.984741. Erratum in: Front. Microbiol. 14, 1279422. doi: 10.3389/fmicb.2023.1279422.

Jabar, A.K., Romli, N.IA., Vellasamy, K.M., Pallath, V., Al-Maleki, A.R. (2026) Predictors of Mortality in Pseudomonas aeruginosa Bloodstream Infections: A Scoping Review. Pathogens 15(1), 61. doi: 10.3390/pathogens15010061.

Kapoor, R., Selvaraju, S.B., Subramanian, V., Yadav, J.S. (2024) Microbial Community Establishment, Succession, and Temporal Dynamics in an Industrial Semi-Synthetic Metalworking Fluid Operation: A 50-Week Real-Time Tracking. Microorganisms 12, 267. 10.3390/microorganisms12020267.

Khan, I.U., Selvaraju, S.B., Yadav, J.S. (2005) Occurrence and characterization of multiple novel genotypes of Mycobacterium immunogenum and Mycobacterium chelonae in metalworking fluids. FEMS Microbiol. Ecol. 54(3), 329–338. doi: 10.1016/j.femsec.2005.04.009.

Koch, T., Passman, F., Rabenstein, A. (2015) Comparative study of microbiological monitoring of water-miscible metalworking fluids. Int. Biodeterior. Biodegrad. 98, 19–25. 10.1016/j.ibiod.2014.11.015.

Koch, M., Rosiwal, S., Burkovski, A. (2022) Environmentally sustainable elimination of microbes using boron-doped diamond electrodes: From water treatment to medical applications. In: Chowdhary, P., Mani, S., Chaturvedi, P. (eds.) Microbial biotechnology: Role in ecological sustainability and research. John Wiley & Sons, Inc., 355–364.

Kraft, A. (2007) Doped diamond: A compact review on a new, versatile electrode material. Int. J. Electrochem. Sci. 2, 355–385.

Kreiss, K., Cox-Ganser, J. (1997) Metalworking fluid-associated hypersensitivity pneumonitis: a workshop summary. Am. J. Ind. Med. 32(4), 423–432. doi: 10.1002/(sici)1097-0274(199710)32:4<423::aid-ajim16>30.co;2-5.

Leeb-Zatorska, B., Van den Nest, M., Ebner, J., Moser, D., Spettel, K., Bovier-Azula, L., Diab-El Schahawi, M., Presterl, E. (2024) Tolerance of Pseudomonas oleovorans biofilms to disinfectants commonly used in endoscope reprocessing? Biofilm 8, 100221. doi: 10.1016/j.bioflm.2024.100221.

Lefèvre, C.R., Pelletier, R., Le Monnier, A., Corvec, S., Bille, E., Potron, A., Fihman, V., Farfour, E., Amara, M., Degand, N., Barraud, O., Cattoir, V., for The Gmc Study Group. (2021) Clinical relevance and antimicrobial susceptibility profile of the unknown human pathogen Corynebacterium aurimucosum. J. Med. Microbiol. 70(3). doi: 10.1099/jmm.0.001334.

Liu, B.M., Beck, E.M., Fisher, M.A. (2021a) The Brief Case: Ventilator-Associated Corynebacterium accolens Pneumonia in a Patient with Respiratory Failure Due to COVID-19. J. Clin. Microbiol. 59(9), e0013721. doi: 10.1128/JCM.00137-21.

Liu, B.M., Beck, E.M., Fisher, M.A. (2021b) Closing the Brief Case: Ventilator-Associated Corynebacterium accolens Pneumonia in a Patient with Respiratory Failure Due to COVID-19. J. Clin. Microbiol. 59(9), e0013821. doi: 10.1128/JCM.00138-21.

Lodders, N., Kämpfer, P. (2012). A combined cultivation and cultivation-independent approach shows high bacterial diversity in water-miscible metalworking fluids. Syst. Appl. Microbiol. 35(4), 246–252. doi: 10.1016/j.syapm.2012.03.006.

Marchand, G., Lavoie, J., Racine, L., Lacombe, N., Cloutier, Y., Bélanger, E., Lemelin, C., Desroches, J. (2010) Evaluation of bacterial contamination and control methods in soluble metalworking fluids. J. Occup. Environ. Hyg. 7(6), 358–366. doi: 10.1080/15459621003741631.

Martirosyan, V., Moosavi, E., Ayrapetyan, S. (2011) The study of the effects of carbon dioxide-induced elevation of hydrogen peroxide toxicity on microbes as a novel tool for water purification. World J. Microbiol. Biotechnol. 27, 1091–1098.

Mattsby-Baltzer, I., Sandin, M., Ahlström, B., Allenmark, S., Edebo, M., Falsen, E., Pedersen, K., Rodin, N., Thompson, R.A., Edebo, L. (1989) Microbial growth and accumulation in industrial metal-working fluids. Appl. Environ. Microbiol. 55(10), 2681–2689. doi: 10.1128/aem.55.10.2681-2689.1989.

Mayser, P. (2025) Scopulariopsis brevicaulis: Der „Arsenpilz” – ein schwierig zu behandelnder Erreger [Scopulariopsis brevicaulis: The “arsenic fungus”-a difficult-to-treat pathogen]. Dermatologie (Heidelb). 2025 Sep;76(9):533–543. German. doi: 10.1007/s00105-025-05526-9.

Menberu, M.A., Liu, S., Cooksley, C., Hayes, A.J., Psaltis, A.J., Wormald, P.J., Vreugde, S. (2021) Corynebacterium accolens Has Antimicrobial Activity against Staphylococcus aureus and Methicillin-Resistant S. aureus Pathogens Isolated from the Sinonasal Niche of Chronic Rhinosinusitis Patients. Pathogens 10(2), 207. doi: 10.3390/pathogens10020207.

Moore, J.S., Christensen, M., Wilson, R.W., Wallace, R.J. Jr., Zhang, Y., Nash, D.R., Shelton, B. (2000) Mycobacterial contamination of metalworking fluids: involvement of a possible new taxon of rapidly growing mycobacteria. AIHAJ. 61(2), 205–213. doi: 10.1080/15298660008984529.

Murat, J.-B., Grenouillet, F., Reboux, G., Penven, E., Batchili, A., Dalphin, J.-C., Thaon, I., Millon, L. (2012) Factors Influencing the Microbial Composition of Metalworking Fluids and Potential Implications for Machine Operator’s Lung. Appl. Environ. Microbiol. 78, 34–41. 10.1128/AEM.06230-11.

Natsis, N.E., Cohen, P.R. (2018) Coagulase-Negative Staphylococcus Skin and Soft Tissue Infections. Am. J. Clin. Dermatol. 19(5), 671–677. doi: 10.1007/s40257-018-0362-9.

Núñez-García, L.Á., Córdova-Fletes, C., Barboza-Cerda, M.C., Garza-González, E. (2026) Pseudomonas aeruginosa Biofilms in Cystic Fibrosis: Interactions, Methods, and Therapeutic Strategies. Biomed. Res. Int. 2026, 5328382. doi: 10.1155/bmri/5328382.

O’Flaherty, E., Membré, J.-M., Cummins, E. (2018) Meta-analysis of the reduction of sensitive and antibiotic-resistant Escherichia coli as a result of low and medium pressure UV lamps. Water Sci. Technol. DOI: 10.2166/wst.2018.183.

Parekh, V.R., Traxler, R.W., Sobek, J.M. (1977) N-Alkane oxidation enzymes of a pseudomonad. Appl. Environ. Microbiol. 33, 881–884. 10.1128/aem.33.4.881-884.1977.

Passman, F.J., Küenzi, P. (2020) Microbiology in Water-Miscible Metalworking Fluids. Tribology Trans. 63(6), 1147-1171. Doi.org/10.1080/10402004.2020.1764684.

Perkins, S.D., Angenent, L.T. (2010) Potential pathogenic bacteria in metalworking fluids and aerosols from a machining facility: Potential pathogens in metalworking fluids and aerosols. FEMS Microbiol. Ecol. 74, 643–654. 10.1111/j.1574-6941.2010.00976.x.

Poucke, L.V., Hinton, J.B., Heck, H.C., Heck, B.E., Heck, B.E. (2025) Systemic Sclerosis Presenting as Osteomyelitis of the Finger: Physicians Must Maintain a High Index of Suspicion for Systemic Sclerosis when Evaluating Patients with Fingertip Ulceration or Infection. J. Orthop. Case Rep. 15(4), 52–55. doi: 10.13107/jocr.2025.v15.i04.5442.

Rabenstein, A., Koch, T., Remesch, M., Brinksmeier, E., Kuever, J. (2009) Microbial degradation of water miscible metal working fluids. Int. Biodeterior. Biodegrad. 63, 1023–1029. 10.1016/j.ibiod.2009.07.005.

Ravaioli, S., De Donno, A., Bottau, G., Campoccia, D., Maso, A., Dolzani, P., Balaji, P., Pegreffi, F., Daglia, M., Arciola, C.R. 2024. The Opportunistic Pathogen Staphylococcus warneri: Virulence and Antibiotic Resistance, Clinical Features, Association with Orthopedic Implants and Other Medical Devices, and a Glance at Industrial Applications. Antibiotics 13(10), 972. doi: 10.3390/antibiotics13100972.

Reeve, M.A., Buddie, A.G. (2018) A simple and inexpensive method for practical storage of field-sample proteins for subsequent MALDI-TOF MS analysis. Plant Methods 14, 90. doi: 10.1186/s13007-018-0358-8.

Rosenman, K.D. (2009) Asthma, hypersensitivity pneumonitis and other respiratory diseases caused by metalworking fluids. Curr. Opin. Allergy Clin. Immunol. 9(2), 97–102. doi: 10.1097/ACI.0b013e3283229f96.

Saha, R., Donofrio, R.S. (2012) The microbiology of metalworking fluids. Appl. Microbiol. Biotechnol. 94(5), 1119–1130. doi: 10.1007/s00253-012-4055-7.

Santos, D.K., Rufino, R.D., Luna, J.M., Santos, V.A., Sarubbo, L.A. (2016) Biosurfactants: Multifunctional Biomolecules of the 21st Century. Int. J. Mol. Sci. 17(3), 401. doi: 10.3390/ijms17030401.

Schorr, B., Ghanem, H., Rosiwal, S., Geißdörfer, W., Burkovski, A. (2019) Elimination of bacterial contaminations by treatment of water with diamond electrodes. World J. Microbiol. Biotechnol. 35, 48.

Schwermer, C.U., Krzeminski, P., Wennberg, A.I., Vogelsang, C., Uhl, W. (2018) Removal of antibiotic-resistant E. coli in two Norwegian wastewater treatment plants and by nano- and ultrafiltration processes. Water Sci. Technol. 77, 1115–1126.

Shelton, B.G., Flanders, W.D., Morris, G.K. (1999) Mycobacterium sp. as a possible cause of hypersensitivity pneumonitis in machine workers. Emerg. Infect. Dis. 5(2), 270–273. doi: 10.3201/eid0502.990213.

Sloyer, J.L., Novitsky, T.J., Nugent, S. (2002) Rapid bacterial counts in metal working fluids. J. Ind. Microbiol. Biotechnol. 29(6), 323–324. doi: 10.1038/sj.jim.7000305.

Theaker, D., Thompson, I. (2010) The Industrial Consequences of Microbial Deterioration of Metal-Working Fluid, in: Timmis, K.N. (Ed.), Handbook of Hydrocarbon and Lipid Microbiology. Springer Berlin Heidelberg, Berlin, Heidelberg, pp. 2641–2650. 10.1007/978-3-540-77587-4_196.

Thorne, P.S., Adamcakova-Dodd, A., Kelly, K.M., O’neill, M.E., Duchaine, C. (2006) Metalworking fluid with mycobacteria and endotoxin induces hypersensitivity pneumonitis in mice. Am. J. Respir. Crit. Care Med. 173(7), 759–768. doi: 10.1164/rccm.200405-627OC.

Tillie-Leblond, I., Grenouillet, F., Reboux, G., Roussel, S., Chouraki, B., Lorthois, C., Dalphin, J.C., Wallaert, B., Millon, L. (2011) Hypersensitivity pneumonitis and metalworking fluids contaminated by mycobacteria. Eur. Respir. J. 37(3), 640–647. doi: 10.1183/09031936.00195009.

Topić Popović, N., Kazazić, S.P., Bojanić, K., Strunjak-Perović, I., Čož-Rakovac, R. (2023) Sample preparation and culture condition effects on MALDI-TOF MS identification of bacteria: A review. Mass Spectrom. Rev. 42(5), 1589–1603. doi: 10.1002/mas.21739.

Toribio, J., Escalante, A.E., Soberón-Chávez, G. (2010) Production of rhamnolipids in bacteria other than Pseudomonas aeruginosa. Eur. J. Lipid Sci. Technol. 112, 1082–1087.

Trafny, E.A., Lewandowski, R., Kozłowska, K., Zawistowska-Marciniak, I., Stępińska, M. (2015) Microbial contamination and biofilms on machines of metal industry using metalworking fluids with or without biocides. Int. Biodeterior. Biodegrad. 99, 31–38. 10.1016/j.ibiod.2014.12.015.

van Beilen, J.B., Funhoff, E.G. (2007) Alkane hydroxylases involved in microbial alkane degradation. Appl. Microbiol. Biotechnol. 74(1), 13–21. doi: 10.1007/s00253-006-0748-0.

van Beilen, J.B., Wubbolts, M.G., Witholt, B. (1994) Genetics of alkane oxidation by Pseudomonas oleovorans. Biodegradation 5(3-4), 161–174. doi: 10.1007/BF00696457.

van der Gast, C.J., Whiteley, A.S., Lilley, A.K., Knowles, C.J., Thompson, I.P. (2003) Bacterial community structure and function in a metal-working fluid. Environ. Microbiol. 5(6), 453–461. doi: 10.1046/j.1462-2920.2003.00428.x.

Veillette, M., Thorne, P., Gordon, T. Duchaine, C. (2004) Six Month Tracking of Microbial Growth in a Metalworking Fluid After System Cleaning and Recharging. Ann. Occup. Hyg. 48 (6), 541–546. 10.1093/annhyg/meh043.

Wallace, R.J. Jr., Zhang, Y., Wilson, R.W., Mann, L., Rossmoore, H. (2002) Presence of a single genotype of the newly described species Mycobacterium immunogenum in industrial metalworking fluids associated with hypersensitivity pneumonitis. Appl. Environ. Microbiol. 68(11), 5580–5584. doi: 10.1128/AEM.68.11.5580-5584.2002.

Weimer, P.J. (2022) Degradation of Cellulose and Hemicellulose by Ruminal Microorganisms. Microorganisms 10(12), 2345. doi: 10.3390/microorganisms10122345.

Wilson, R.W., Steingrube, V.A., Böttger, E.C., Springer, B., Brown-Elliott, B.A., Vincent, V., Jost, K.C., Zhang, Y., Garcia, M.J., Chiu, S.H., Onyi, G.O., Rossmoore, H., Nash, D.R., Wallace, R.J. (2001) Mycobacterium immunogenum sp. nov., a novel species related to Mycobacterium abscessus and associated with clinical disease, pseudo-outbreaks and contaminated metalworking fluids: an international cooperative study on mycobacterial taxonomy. Int. J. Syst. Evol. Microbiol. 51(Pt 5), 1751–1764. doi: 10.1099/00207713-51-5-1751.

Wong, J.S., Seaward, L.M., Ho, C.P., Anderson, T.P., Lau, E.O., Amodeo, M.R., Metcalf, S.C., Pithie, A.D., Murdoch, D.R. (2010) Corynebacterium accolens-associated pelvic osteomyelitis. J. Clin. Microbiol. 48(2), 654–655. doi: 10.1128/JCM.00818-09.

Zacharisen, M.C., Kadambi, A.R., Schlueter, D.P., Kurup, V.P., Shack, J.B., Fox, J.L., Anderson, H.A., Fink, J.N. (1998) The Spectrum of Respiratory Disease Associated With Exposure to Metal Working Fluids. J. Occup. Environ. Med. 40, 640. 10.1097/00043764-199807000-00010.

